# Generation of miR-141/200c conditional knockout mice from knockout-first, reporter-tagged parent and functional validation of the floxed allele

**DOI:** 10.1101/2025.09.02.673774

**Authors:** Sanjeev Kumar Yadav, Rashmi Srivastava, Mary-Katherine Cormier, Katie Lowther, Siu-Pok Yee, Rajkumar Verma

## Abstract

MicroRNAs (miRNAs) of the miR-200 family—specifically miR-141 and miR-200c— regulate neurogenesis, differentiation, and epithelial–mesenchymal transitions in development. Dysregulation of these miRNAs is associated with several diseases including cancer and stroke. The Mirc13tm1Mtm/Mmjax mouse line, which targets the miR-141/200c cluster, was originally generated and described by Park et al. 2012 as a knockout-first, reporter-tagged insertion with conditional potential (conditional-ready) mouse line. Harnessing its full potential requires a two-step breeding process: breeding with FLP mice to excise the lacZ/neo cassette, then breeding with Cre to delete the floxed miRNA cluster (Park, et al. 2012). However, many studies either bypassed removal of the lacZ/Neo cassettes and treated the mouse line as Mirc13 knockouts or bred directly with Cre mouse lines, which could lead to unpredictable recombination and genotypes. Here we show that retention of the lacZ/Neo cassette is associated with reduced expression of the neighboring genes Ptpn6, Phb2 and Atn1 in the olfactory bulb, and that these genes are expressed normally once the cassette is excised. We therefore recommend a validated two-step FLPo-then-Cre breeding plan for this line, together with case-by-case allele validation for other knockout-first, reporter-tagged mouse lines.

## INTRODUCTION

microRNAs (miRNAs) are small, noncoding RNAs (∼22 nucleotides) that regulate gene expression post-transcriptionally and play pivotal roles in development and disease. Originally discovered in C. elegans (Lee, et al. 1993; Wightman, et al. 1993) and later shown to be evolutionarily conserved across species (Pasquinelli, et al. 2000), miRNAs are now recognized as critical regulators in cardiovascular (Small, et al. 2010; van Rooij and Kauppinen 2014), neurological (Wu, et al. 2016; Verma, et al. 2018), and oncogenic pathways (Humphries and Yang 2015; Jin, et al. 2017), holding significant potential for therapeutic applications (Janssen, et al. 2013; Verma, et al. 2018). While cell-based studies have elucidated many of these mechanisms, *in vivo* models remain essential to fully understand the functions of miRNAs under physiological and pathological contexts. Park et al. established a key genetic resource for *in vivo* miRNA investigation by generating conditional knockout-first mouse lines for conserved miRNAs, including the miR-141/200c cluster (Mirc13tm1Mtm/Mmjax) (Park, et al. 2012). This cluster, located on mouse chromosome 6, encodes two members of the miR-200 family implicated in neurogenesis (Choi, et al. 2008; Beclin, et al. 2016), differentiation (Zhang, et al. 2015), epithelial–mesenchymal transition (Tseng, et al. 2017), cancer (Chen, et al. 2013; Jin, et al. 2017), stroke (Small, et al. 2010; Yadav, et al. 2024), and Alzheimer’s disease (Wu, et al. 2016). To generate miR-141/200c cluster KO mice, Park et al. (2012) used a targeting vector to incorporate a lacZ reporter and β-actin-neo selection cassette flanked by FRT sites and loxP sites surrounding the miR-141/200c cluster. This design was aimed towards a two-step recombination strategy: first, FLP-mediated removal of the lacZ/neo cassette generates a classical conditional (floxed) allele, and then Cre-mediated excision to remove the miR-141/200c cluster (Park, et al. 2012). However, many subsequent studies bypassed the recommended FLP recombination step (Guan, et al. 2018; Hoefert, et al. 2018; Ji, et al. 2018; Liu, et al. 2018; Wu, et al. 2019; Xu, et al. 2025). In such cases, the retained lacZ/neo selection cassette could alter neighboring gene expression through multiple potential mechanisms, including promoter-mediated transcriptional interference or disruption of local transcriptional organization, and/or imposing an unanticipated neighborhood effect (Pham, et al. 1996) thereby confounding phenotypic interpretation (Testa, et al. 2004; Raja, et al. 2024). In few other studies attempted cassette removal using FLPe deleter strains; however, several commonly used FLPe lines exhibit variable recombination efficiency, which may lead to mosaic excision depending on the deleter strain and genomic target locus (Tran, et al. 2017; Mostofa, et al. 2022; Tran, et al. 2024). FLPe-mediated excision can therefore produce mosaic founder animals. Founder mosaicism is not, however, an inherent limitation of FLPe and does not propagate as cohort heterogeneity: an additional backcross to wild-type, with genotype selection and segregation of the recombinase allele, routinely yields offspring carrying a defined, fully excised allele. A codon-optimized FLPo deleter reduces the number of founders that must be screened, but is not a prerequisite for obtaining a clean allele. Several studies have reported that residual selection cassettes can unintentionally influence expression of nearby genes (Olson, et al. 1996; Pham, et al. 1996; Skarnes, et al. 2011). In the case of the miR-141/200c cluster, expression of genes located in close proximity (www.Ensembl.org) including *Ptpn6* (Gene ID 15170)*, Phb2* (Gene ID 12034)*, Atn1* (Gene ID 13498)*, Eno2* (Gene ID 13807) and *Emg1* (Gene ID 14791) may be affected. These five genes were selected on two pre-specified criteria: each lies within 50 kb of the Mirc13 insertion site, the interval over which selection-cassette effects on neighboring transcription are most often reported, and each has an established role in neuronal function, neurodevelopment, mitochondrial integrity or ribosomal biogenesis, so that altered expression would plausibly confound a neural phenotype (Whitton, et al. 1993; Eschrich, et al. 2002; Wu, et al. 2010; Xu, et al. 2014; Isgrò, et al. 2015; He, et al. 2023), altered expression of these genes could lead to misleading interpretations or the apparent loss of phenotype in knockout mice generated due to the presence of the neo cassette with a strong promoter (Table 1).

**Table 1.** Summary of published miR-141/200c knockout models and the potential influence of neighboring genes on reported phenotypes.

| Study | Methodology/mouse model | Recombinase Used | Reported miR-141/200c Phenotype | Neighboring Gene(s) Potentially Affecting Phenotype | Evidence from Independent Gene Deletion/Functional Studies |
| --- | --- | --- | --- | --- | --- |
| Guan et al., 2018 | Conditional miR-200c/141 floxed mice crossed with CD4-Cre mice to generate T-cell-specific miR-141/200c deletion | CD4-Cre (no information provided about crossing with FLP) | Increased terminal effector CD8<sup>+</sup> T cells, reduced memory T-cell formation, altered ZEB1/ZEB2 network | Ptpn6 (High potential) | PTPN6/SHP-1 is a critical negative regulator of T-cell receptor signaling (Lorenz 2009). |
| Hoefert et al., 2018. | miR-141/200c heterozygous mice crossed with K14-Cre | K14-Cre | miR-200 loss disrupted epithelial adhesion, proliferation, and hair morphogenesis. | No major candidate neighboring gene identified | Not applicable |
| Ji et al., 2018. | miR-141/200c heterozygous mice were intercrossed to generate homozygous Mirc13 knockout mice | No information provided. | Reduced disc degeneration, apoptosis, ECM degradation | Phb2 (Moderate potential) | PHB2 loss impairs mitochondrial function, increases oxidative stress, and promotes cell injury (Merkwirth and Langer 2009). |
| Liu et al., 2018. | miR-141/200c floxed mice crossed with mammary epithelial Cre line | MMTV-Cre (no information provided about crossing with FLP) | miR-141/200c regulated breast cancer stem cell heterogeneity through HIPK1/β-catenin signaling. | No major candidate neighboring gene identified | Not applicable |
| Wu et al., 2016. | miR-141/200c heterozygous mice crossed with mammary epithelial Cre line | MMTV-Cre (no information provided about crossing with FLP) | Mammary-specific miR-200c deletion disrupted epithelial polarity, promoted EMT, and altered stem cell behavior by affecting mitochondrial dynamics. | Phb2 (Moderate potential) | PHB2 is a mitochondrial inner membrane protein involved in mitochondrial organization, apoptosis regulation, and PINK1/Parkin-dependent mitophagy (Yan, et al. 2020). |
| Xu et al., 2025. | miR-141/200c heterozygous mice crossed with the tamoxifen-inducible Cre-recombinase strain B6;129S6-Tg, CKIIα-cre/ERT2 | tamoxifen-inducible Cre-recombinase strain B6;129S6-Tg, CKIIα-cre/ERT2 | Neuronal miR-141/200c deletion reduced ischemic injury and improved stroke outcomes. | Phb2, Atn1, Eno2 (Moderate potential) | PHB2 regulates mitochondrial homeostasis and is important for neuronal survival (Yan, et al. 2020). ATN1 regulates neuronal transcription and its mutation causes neurodegenerative phenotypes. |

The two-step FLP-then-Cre scheme is established best practice for knockout-first alleles, and we do not present it as a methodological advance. Our contribution is the experimental validation of genetic integrity in a widely used line. Using a codon-optimized ROSA26-FLPo deleter (Raymond and Soriano 2007; Wu, et al. 2009; Kranz, et al. 2010), we excised the lacZ/neo cassette to generate a conventional floxed allele before Cre-mediated deletion, and we compared neighboring-gene expression before and after excision. The complete allele series and the breeding steps used to generate it are summarized in Figure 1, and we report the genotyping assays needed to distinguish every intermediate allele (Table 2, Figures 1 and 6). The value of these data lies in demonstrating what is lost when a standard validation step is omitted, and in providing a validated breeding and genotyping framework for generating unambiguous global or cell-specific miR-141/200c knockouts.

**Fig 1.**
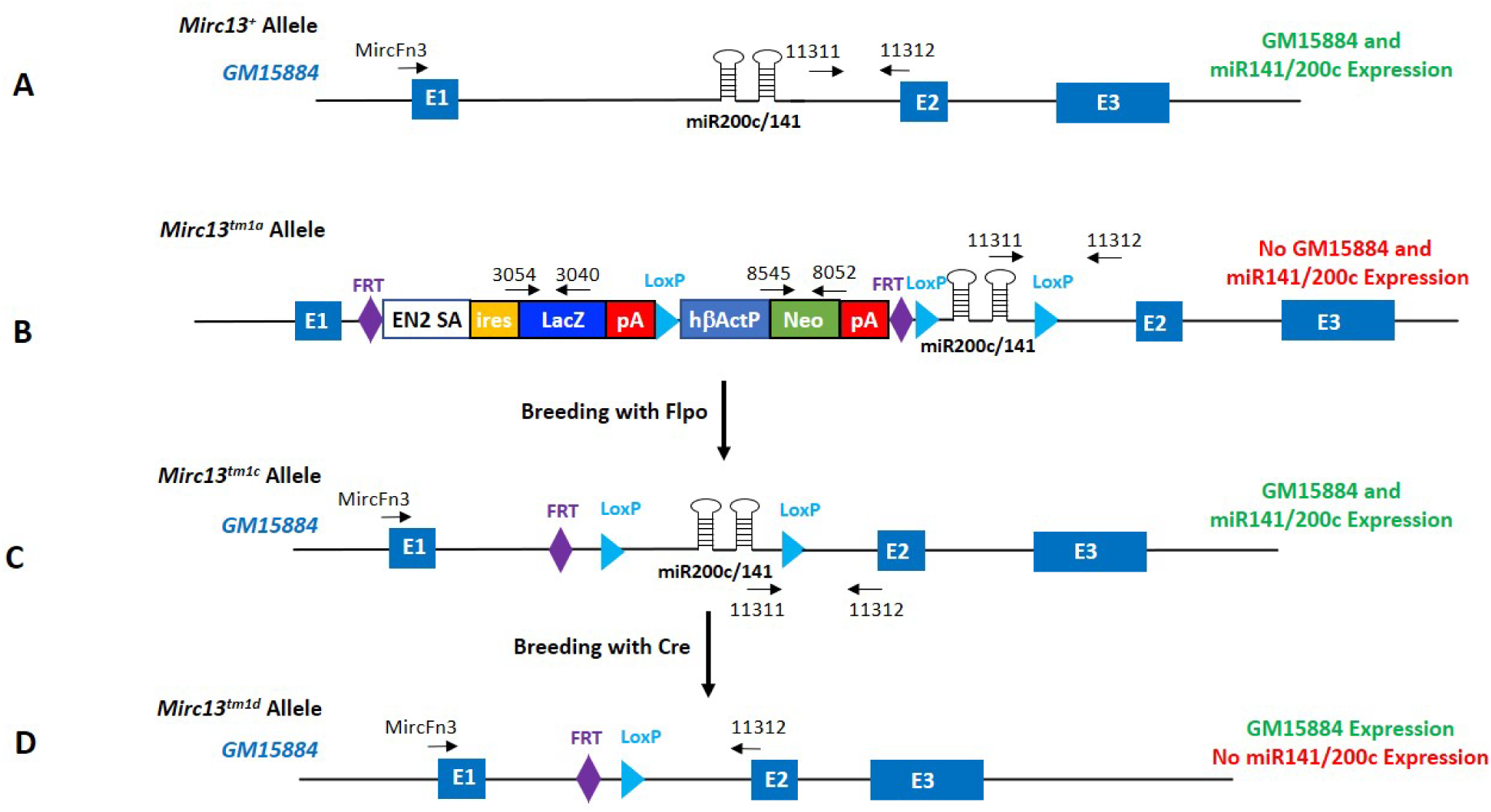
Genomic organization of *Mirc13*, *Mirc13t^m1a^,* after Flpo and Cre mediated recombination. A. *Mirc13*, consists of *miR-200/144c cluster*, is located in intron 1 of lncRNA, *GM15884*; B. The *Mirc13^tm1a^* mouse line was originally designed as a knockout-first, reporter-tagged insertion with conditional potential (conditional-ready) mouse line for *Mirc13* (*miR-200c/141* cluster). Insertion of the En2-lacZ gene trap cassette was designed to prevent subsequent 3’-downstream transcription that could lead to the lack of *GM15884* and *Mirc13* expressions. C. When these mice were bred with *Rosa26-Flpo*, *Flpo* will facilitate excision of the LacZ/Neo cassette to generate the *Mirc13^tm1c^* allele with wildtype level of expression of both *GM15884* and *miR-200c/141*. D. When *Mirc13^tm1c^* mice were bred with Cre, Cre will mediate excision of the *Mirc13* cluster sequence to generate *Mirc13^tm1d^*allele with wildtype *GM15884* expression and the lack of *miR-200c/141* expression.

**Table 2.**
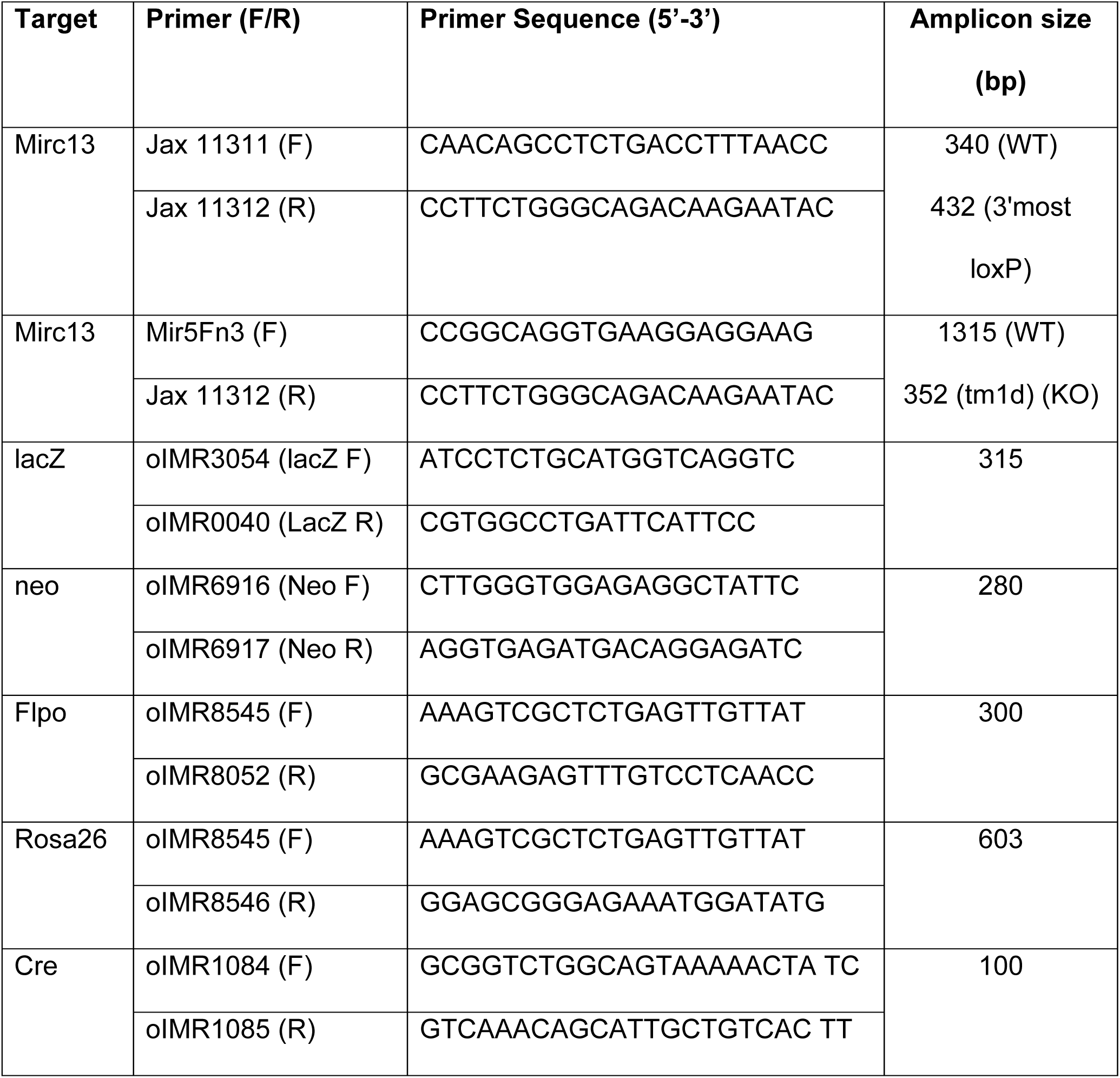
List of different sets of primer sequences (with amplicon size) used in PCR amplification. The corresponding position of the primers is shown in Figure 1 and 6.

## MATERIAL AND METHODS

### Mouse Strains

All mouse strains used in this study were obtained from the Jackson Laboratory (Bar Harbor, ME): Mirc13^tm1Mtm^/Mmjax line (MMRRC stock #034657 (Park, et al. 2012)) (hereon referred as *Mirc13^tm1^* line); B6.129S4-Gt(ROSA)26Sor^tFLP^)Sor/J (ROSA-FLPo; Jax# 012930) (Raymond and Soriano 2007); 129S1/Sv-Hprt1^tm1(CAG-cre)Mnn^/J) (Hprt-Cre; Jax# 004302) (Tang, et al. 2002) and B6.129P2-Lyz2^tm1(cre)Ifo^/J) (LysMCre; Jax # 004781) (Clausen, et al. 1999). All mice were maintained under standard conditions at ambient temperature and humidity, with ad libitum access to pellet food and water. All animal studies were approved by the Institutional Animal Care and Use Committee at the University of Connecticut Health Center. Unless stated otherwise, expression analyses used n = 3 mice per genotype (2 females, 1 male) at 3-4 weeks of age, with each sample assayed in technical duplicate; sex was recorded, but the study was not powered to test sex as a biological variable. Parent of origin of the targeted allele was as follows: Mirc13tm1a/tm1a mice were generated by intercrossing male and female Mirc13tm1a/+ animals, whereas the Mirc13tm1c and Mirc13tm1d lines were generated from female Mirc13tm1a/+ mice crossed with male Rosa26FLPo/FLPo mice, and the resulting Mirc13tm1c/+ animals were crossed with Hprttm1(CAG-cre)Mnn mice.

### Genotyping of Mouse Strains

Tail biopsies (∼4 mm) were collected in DNase-free microtubes and incubated in 50 µl of alkali buffer (25mM NaOH and 0.2mM EDTA) at 100°C for one hour and an equal volume of neutralization buffer (40mM Tris buffer, pH 7.5) was added. One microliter of the DNA was used for PCR genotyping with 2× JumpStart™ Taq ReadyMix (Sigma-Aldrich P2893). Primer pairs are listed in Table 2. PCR was performed according to the manufacturer’s instructions. PCR products were fractionated on 2% agarose gels prepared in 1× Tris-Acetate-EDTA (TAE) buffer (Bio-Rad 1610743) with SYBR™ Safe DNA stain (Invitrogen S33102) and visualized under UV illumination.

### β-Galactosidase (lacZ) Staining

Mice were deeply anesthetized with Avertin (250 mg/kg, i.p.), perfused with ice-cold PBS, and brains were collected for cryosections (10 µm, Superfrost Plus slides). Sections were fixed with fixation solution (provided with kit) for 10 min, and LacZ expression was determined using a β-galactosidase reporter staining kit (Sigma-Aldrich, GALS-1KT) according to manufacturer’s instructions and as in (Burn 2012). Slides were mounted in 70% glycerol and imaged under bright-field microscopy. The specificity and standard use of X-gal staining for detection of beta-galactosidase reporter activity have been described previously (Burn 2012).

### RNA Isolation, cDNA Synthesis, and Real-Time PCR

Total RNA was extracted from the olfactory bulb, frontal and parietal cortex, hippocampus, cerebellum, heart, lung, and liver using TRIzol or miRVana kits (Ambion Life Technologies, 15596018, AM1560). cDNA synthesis for miRNAs and mRNAs was performed using the TaqMan MicroRNA Reverse Transcription Kit (Thermo Fisher 4366596) and iScript cDNA Synthesis Kit (Bio-Rad 1708891), respectively. TaqMan assays included miR-141-3p (000463), miR-200c-3p (002300), U6snRNA (001973), *Ptpn6* (Mm00469153_m1), *Phb2* (Mm00476104_m1), *Atn1* (Mm00492256_m1), *Eno2* (Mm00469062_m1), *Emg1* (Mm00496596_m1), and *Gapdh* (Mm99999915_g1) (Thermo Fisher Scientific). PCR was performed using TaqMan Universal PCR Master Mix II (No UNG; Thermo Fisher 4440040). Relative expression was determined by the 2^–ΔΔCt method using U6 and Gapdh as normalizers for miRNA and mRNA targets, respectively. Fold-Change (2^–ΔΔCt)) is the normalized gene expression (2^(-ΔCt)) in the KO Sample divided the normalized gene expression (2^(-ΔCt)) in the WT. The p values are calculated based on a Student’s t-test of the replicate 2^(-ΔCt) values for each gene in the WT group and KO groups, and p<0.05 are significant. Statistical testing is described under Statistical Analysis.

### RNA Fluorescence In Situ Hybridization (RNA-FISH)

RNA-FISH was performed using Molecular Instruments protocols. Fixed tissue sections were pre-hybridized at 37 °C and incubated overnight with 16 nM of the miR-141-3p probe. Following stringent SSCT washes, signal amplification was performed using snap-cooled hairpins (h1/h2; 6 pmol each) incubated overnight at room temperature. Sections were mounted with antifade DAPI UltraCruz Mounting Solution (Santa Cruz SC-24941) and imaged by fluorescence microscopy.

### Statistical Analysis

Data are presented as mean ± SD. Normalized expression values derived from the 2^– ΔCt method were used for statistical analysis. Statistical analyses were performed using GraphPad Prism (San Diego, CA). Distributional assumptions were assessed by the Shapiro–Wilk test. Data met the normality assumption except where the knockout group lay at the detection floor with near-zero variance (miR-141-3p and miR-200c-3p in cassette-carrying and null tissue); these comparisons represent essentially complete loss of signal and are supported by RNA-FISH in addition to RT-qPCR. Given the group size (n = 3), we note that this test has limited power and does not treat a non-significant result as evidence of normality. Comparisons between two groups used a two-tailed unpaired Student’s t-test and multiple-group comparisons one-way ANOVA with Bonferroni post hoc correction; non-parametric equivalents (Mann–Whitney U, Kruskal–Wallis) were performed as a consistency check and gave concordant conclusions, with the caveat that at n = 3 per group the exact two-sided Mann–Whitney p value cannot fall below 0.10 (so data was not shown). Exact p values are reported in each figure, and each dot represents an individual mouse. Blinding was implemented during data acquisition and analysis where feasible.

## RESULTS

### Genomic Organization of the Mirc13 Mouse Lines and Genotyping of the Mouse Lines

The Mirc13^tm1Mtm^/Mmjax line (MMRRC stock #34657) mouse line, hereon referred as *Mirc13^tm1a^* line, available from the Jax MMRRC is a knockout-first, reporter-tagged insertion with conditional potential (conditional-ready) mouse line. We used the Infrafrontier/EMMA allele nomenclature (Skarnes, et al. 1992) from hereon. The miR-141/200c (*Mirc13*) cluster is located within intron 1 of the lncRNA, *GM15884*. As shown in Fig 1B, the targeted allele contains an *engrailed 2* (En2) splice acceptor followed by an internal ribosomal entry site (IRES) sequence coupled to lacZ reporter sequence and polyadenylation signal sequence. This mouse line was originally generated by gene targeting in embryonic stem (ES) cells via homologous recombination. A β-actin-Neo cassette was placed 3’ of the lacZ polyA sequence in the targeting vector to confer G418 selection to help screen for targeted ES cell clones. The Neo cassette and the lacZ reporter was separated by a single loxP site. The lacZ/Neo sequences were flanked by a pair of FRT sites and inserted into intron 1 of *GM15884* 5’ of the floxed *Mirc13* cluster. The targeting strategy places an En2 splice acceptor–IRES-lacZ cassette followed by a polyadenylation sequence within intron 1 of *GM15884*. This design is intended to terminate downstream transcription and disrupt expression of both *GM15884* and the Mirc13 cluster (knockout-first configuration). However, consistent with previous reports that knockout-first alleles may exhibit residual transcriptional read-through (Lindner, et al. 2021), complete transcriptional termination cannot be assumed solely from the allele design. We first performed PCR genotyping of *Mirc13^tm1a/+^* pups using primer pairs as shown in Table 2. As shown in Fig 2A, we detected amplicon of 340 bp and 432 bp specific to the *Mirc13^+/+^* and *Mirc13^tm1c/+^* allele, respectively. We were also able to amplify fragments of 315 bp and 280 bp specific to lacZ and Neo, respectively, confirming the genotype of the *Mirc13^tm1a/+^*mice we obtained from MMRRC/Jax. We then bred the *Mirc13^tm1a/+^*pups together to obtain *Mirc13^tm1a/tm1a^* pups for expression analyses as described below. PCR genotyping only detected the 432 bp fragment specific to the *Mirc13^tm1a^* allele as well as the lacZ and neo specific amplicons. The absence of the *Mirc13^+/+^* specific amplicon of 340 bp confirmed that these were *Mirc13^tm1a/+^* mice (Fig 2A).

**Fig. 2.**
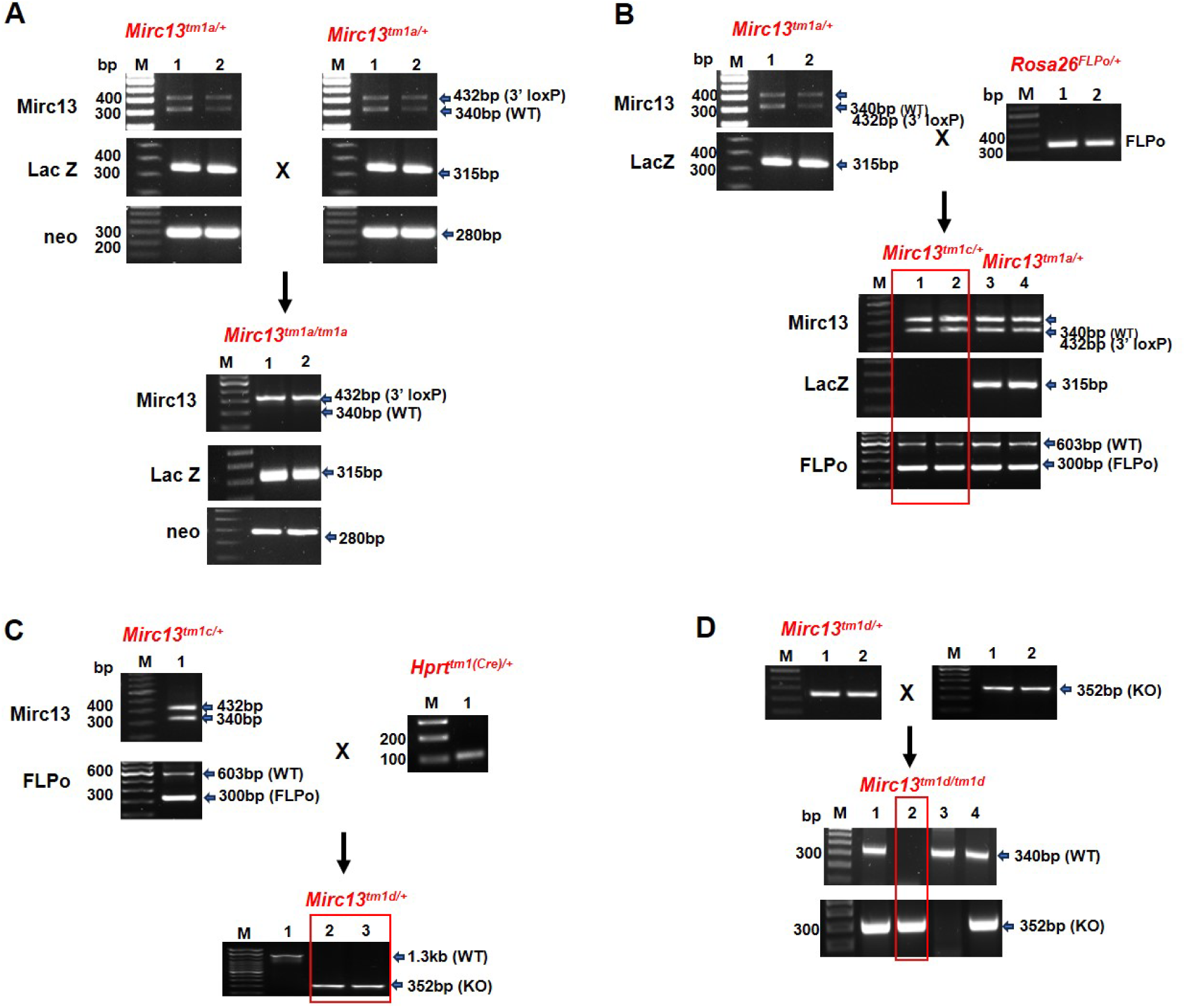
PCR Genotyping of mice with different *Mirc13* alleles. **A** Generation of *Mirc13^tm1a/tm1a^*mice by breeding *Mirc13^tm1a/+^* mice, which contain lacZ/Neo cassette and floxed *Mirc13 sequence*, together; **B** Removal of lacZ and Neo cassette by breeding *Mirc13^tm1a/+^* mice with *Rosa26^FLPo/+^* mice. Positive *Mirc13^tm1c/+^* mice, lacking the lacZ/Neo cassette (315 bp amplicon), were marked by a red box; **C** Generation of *Mirc13^tm1d/+^* (KO) mice by breeding *Mirc13^tm1cl+^* with *Hprt^tm1(Cre)/+^* mice. Positive *Mirc13^tm1d/+^* mice, lacking a 432 bp amplicon specific to the *Mirc13^tm1d/+^* allele, were marked by a red box; **D** *Mirc13^tm1d/+^* mice were bred together to generate *Mirc13^tm1d/tm1d^* global KO mice. These mice were identified by the presence of a KO-specific amplicon (352 bp) and the loss of wt specific amplicon (340 bp) and were marked by the red box. PCR were performed using primer pairs as shown in Table 2 and amplicons specific for lacZ is 315 bp, *Mirc13* is 340 bp, *Mirc13^tm1c^* is 432 bp, *Mirc13^tm1d^* is 352 bp, Rosa26 is 603 bp, Flpo is 300 bp and Cre is 100 bp. Because the JAX-supplied primer pair (Jax 11311/11312) interrogates only the 3′ loxP site, we used a panel of complementary PCRs to assign genotypes unambiguously: lacZ- and Neo-specific reactions to detect the knockout-first cassette, and the Mir5Fn3/Jax 11312 pair spanning the floxed interval to distinguish the wild-type (1.3 kb) and tm1d (352 bp) alleles. Primer positions are shown in Figures 1 and 6 and sequences in Table 2. Under our standard conditions the Mir5Fn3/Jax 11312 reaction preferentially amplifies the shorter tm1d product, so heterozygotes cannot be called from this assay alone; *Mirc13^tm1d/+^* animals are instead identified by the concurrent presence of the tm1d amplicon and the wild-type-specific 340 bp Jax amplicon (Fig. 2D, lanes 1 and 4), whereas *Mirc13^tm1d/tm1d^* mice yield the tm1d amplicon only (Fig. 2D, lane 2). Every animal entering an experimental cohort was genotyped with the full panel.

As described above, the lacZ/Neo sequences are flanked by a pair of FRT sites, which can be excised by Flp recombinase resulting in the generation of functional *Mirc13* conditional knockout (conditional later) mice expressing both *GM15884* and miR-141/200c as shown in Fig 1B. To accomplish this, we bred *Mirc13^tm1a/+^* mice with *Rosa26^Flpo/+^*mice (Jax #012930). As shown in Fig 2B, genotyping results showed the absence of lacZ specific amplicon of 315 bp (lane 1 and 2) thereby confirming successful excision of the lacZ/Neo cassette and generation of *Mirc13^tm1c/+^*pups. To generate *Mirc13^tm1d/tm1d^* (KO) mice, we bred *Mirc13^tm1c/+^*pups with *Hprt^tm1(Cre)/+^* mice (Fig1C). To detect the *Mirc13^tm1d/+^* allele, a forward primer (mir5Fn3) was designed to target the region upstream of the FRT/loxP site and was paired with the Jax 11312 reverse primer downstream of the 3’ loxP site. This primer pair will generate two amplicons, one specific to the *Mirc13^+^* allele (1.3 kb) and one specific to the *Mirc13^tm1d^* allele that is smaller in size due to Cre-mediated recombination (352 bp). As shown in Fig 2C, PCR genotyping showed that we were able to amplify the predicted fragments of 1.3 kb (lane 1) or 352 bp (lanes 2 and 3) specific to *Mirc13^+^* and *Mirc13^tm1d^*alleles, respectively. We anticipated that we should obtain either *Mirc13^+/+^*or *Mirc13^tm1d/+^* pups from this breeding pair. However, we did not detect both amplicons from the *Mirc13^tm1d/+^* mice. This is most likely because our PCR conditions favor amplification of the smaller amplicon specific to the knockout allele, and the pups shown in Fig 2C lanes 2 and 3, were *Mirc13^tm1d/+^* mice. We further bred *Mirc13^tm1d/+^* mice together and genotyped the pups using the same primers as described above for the *Mirc13^tm1d^*allele and also with the original Jax primer pairs specific to the 3’ loxP site in order to detect the presence or absence of the *Mirc13^+^* allele. We were able to obtain *Mirc13^tm1d^*^/tm1d^ mice as shown by PCR genotyping that was positive only for the KO-specific amplicon and no WT-specific amplicon was observed (Fig 2D, lane 2). The other littermates were either *Mirc13^tm1d^*^/+^, which were positive for both amplicons (Fig 2D lanes 1 and 4) or *Mirc13^+/+^* as revealed by the presence of the WT-specific amplicon only (Fig 2D lane 3).

### Expression Analysis of Homozygous Mirc13^tm1^ Mice

The En2 splice acceptor is designed to facilitate splicing between exon 1 of *GM15884* with the IRES-lacZ reporter sequence. Thus, the targeted allele will generate a fusion transcript consisting of exon 1 of the disrupted endogenous *GM15884*, a lncRNA, IRES and lacZ coding sequence. PCR genotyping confirmed presence of the lacZ reporter (Fig 2A). Since the IRES-lacZ gene-trap cassette is inserted into intron 1 of *GM15884* and is under the control of the endogenous *GM15884* promoter, β-galactosidase expression reflects the transcriptional activity of the trapped promoter and serves as a reporter of transcriptional activity at the *GM15884/Mirc13* locus. However, it does not directly reflect the temporal expression pattern or abundance of mature miR-141/200c miRNAs. The knockout-first allele is therefore expected to substantially disrupt expression of both *GM15884* and the *Mirc13* cluster. However, because transcriptional read-through has been reported for some knockout-first alleles (Lindner, et al. 2021), complete transcriptional termination cannot be inferred from the targeting design alone. We first examined lacZ expression, which should reveal endogenous expression of *GM15884* and *miR-141-3p* and *miR-200c-3p*. X-gal staining of the lacZ gene product, β-galactosidase, reflects the transcriptional activity of the endogenous trapped promoter driving *GM15884* expression (Skarnes, et al. 1992), and therefore serves as a surrogate reporter of transcriptional activity at the *GM15884/Mirc13* locus rather than a direct measure of mature miR-141/200c expression. Strong b-galactosidase activity was detected in the olfactory bulb, with weaker staining near the lung–tracheal junction, and minimal expression in the heart and liver (Fig. 3A-C). To further characterize expression of *Mirc13*, we isolated total RNA from the brain olfactory bulb and lungs of *Mirc13^tm1a/tm1a^* and *Mirc13^+/+^* mice and performed RT-qPCR to examine the level of *miR-141-3p* and *miR-200c-3p* expression. Our results showed that *miR-141-3p* and *miR-200c-3p* were highly expressed in the olfactory bulb and lung (ΔCT < 5), whereas heart and liver exhibited lower expression levels (ΔCT > 10) in *Mirc13^+/+^* mice, consistent with the miRNA expression atlas (Rishik, et al. 2025) (Fig. 3C and D). A significant and near-complete reduction of *miR-141-3p* (Fig 3E & F) and *miR-200c-3p* (Fig. 3G & H) expression was observed in the olfactory bulb and lungs in *Mirc13^tm1a/tm1a^*mice. Data are presented as mean ± SD (*p < 0.05 vs. *Mirc13^+/+^*control).

**Fig. 3.**
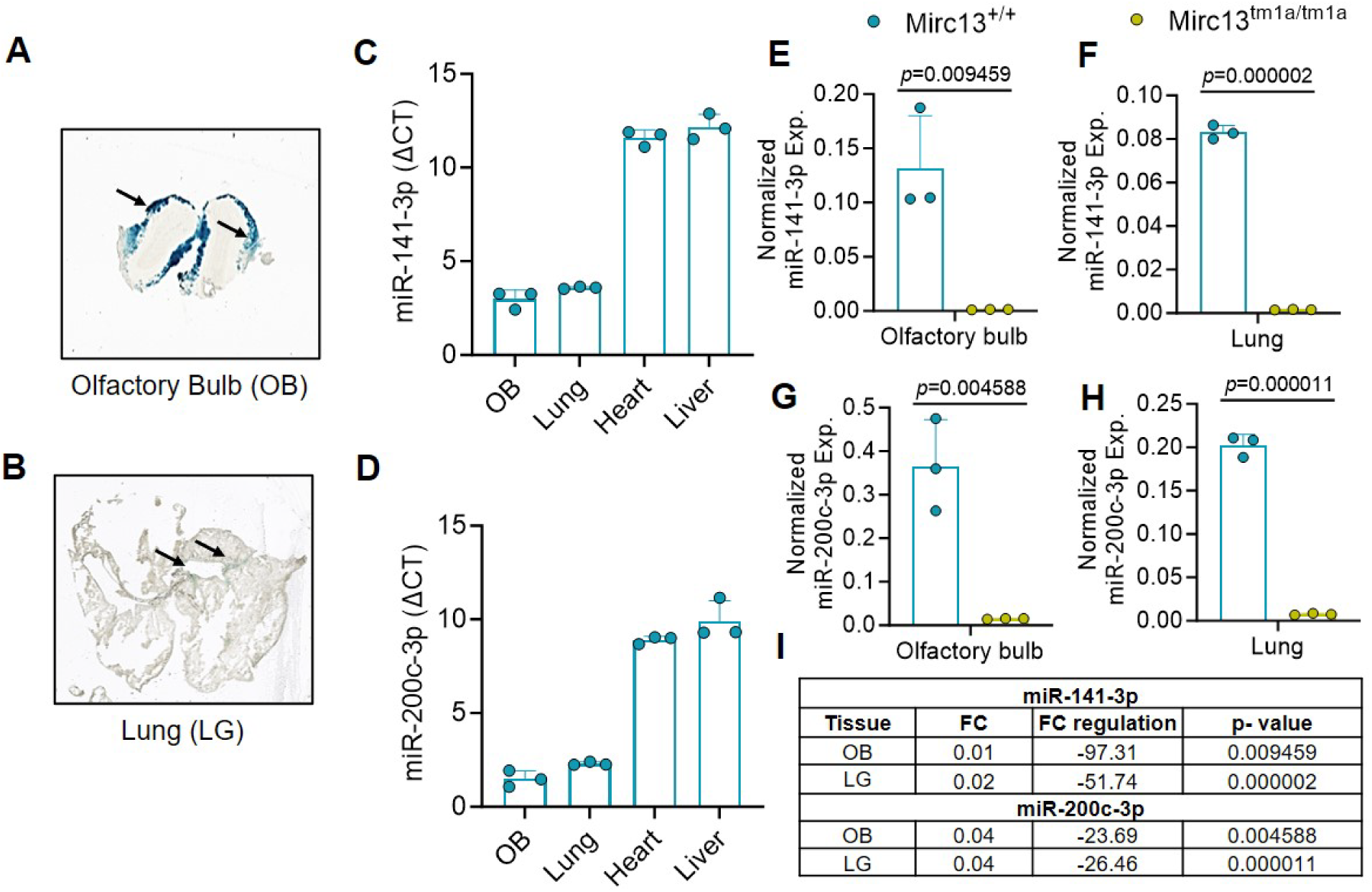
Expression analyses of *Mirc13^tm1a/tm1a^* mice. LacZ expression as revealed by x–gal staining of olfactory bulb (OB) (**A**) and lung (LG) (**B**); Basal expression of *miR-141-3p* and *miR-200c-3p* was analyzed by RT-qPCR in olfactory bulb, lung, heart and liver of wildtype mice (**C**&**D**). Relative expressions of *miR-141-3p* (**E & F**) and *miR-200c-3p* (**G & H**) were assessed by RT-qPCR in the brain olfactory bulb and lungs of *Mirc13^tm1a/tm1a^* mice compared to *Mirc13^+/+^* littermate control mice. Data are presented as mean ± SD (*p < 0.05 vs. *Mirc13^+/+^* littermate control). Blue column denotes *Mirc13^+/+^* and yellow column denotes *Mirc13^tm1a/tm1a^*. Detail Fold Change (FC) regulation of *miR-141-3p* and *miR-200c-3p* (**I**). (n=3 per group, F=2 & M=1, 3-4 Week) each sample run in qPCR with technical duplicate.

### Retention of the lacZ/Neo cassette is associated with reduced neighboring-gene expression

The *Mirc13^tm1a^* mouse line was originally generated by genes targeting in embryonic stem cells via homologous recombination. A lacZ polyA sequence was included in the targeting vector to confer G418 positive selection to screen for targeted ES cell clones. However, the Neo cassette containing a strong ubiquitous β-actin promoter was not removed prior to generation of the *Mirc13^tm1a^* mouse line. It has been reported that insertion of a strong promoter within a gene would not only disrupt endogenous expression, but it could also potentially lead to dysregulation of neighboring genes and resulted in unanticipated phenotypic alterations (Olson, et al. 1996; Pham, et al. 1996; Skarnes, et al. 2011).

To further investigate the influence of the b-actin-Neo cassette on neighboring gene expression, we identified 5 genes, *Ptpn6, Phb2, Atn1, Eno2* and *Emg1* that are important for brain function, located within 50 kb from *Mirc13*. Total RNA was isolated from olfactory bulb of *Mirc13^+/+^*and *Mirc13^tm1a/tm1a^* mice and the levels of neighboring gene expression were characterized by RT-qPCR. As shown in Fig 4A-C, expression of *Ptpn6, Phb2,* and *Atn1* was significantly decreased (±>1.5 folds threshold, p<0.05) in *Mirc13^tm1a/tm1a^* mice as compared to *Mirc13^+/+^* mice, while *Eno2* and *Emg1* showed a similar decreasing trend (Fig 4D and E). In *Mirc13^tm1d/tm1d^* mice, the expression *Ptpn6, Phb2, Atn1, Eno2,* and *Emg1* did not show significantly change (±>1.5 folds threshold, p<0.05) as compared to *Mirc13^tm1c/tm1c^* mice (Fig 4G to K). Retention of the knockout-first cassette is therefore associated with altered expression of neighboring genes; the underlying mechanism is considered in the Discussion.

**Fig. 4.**
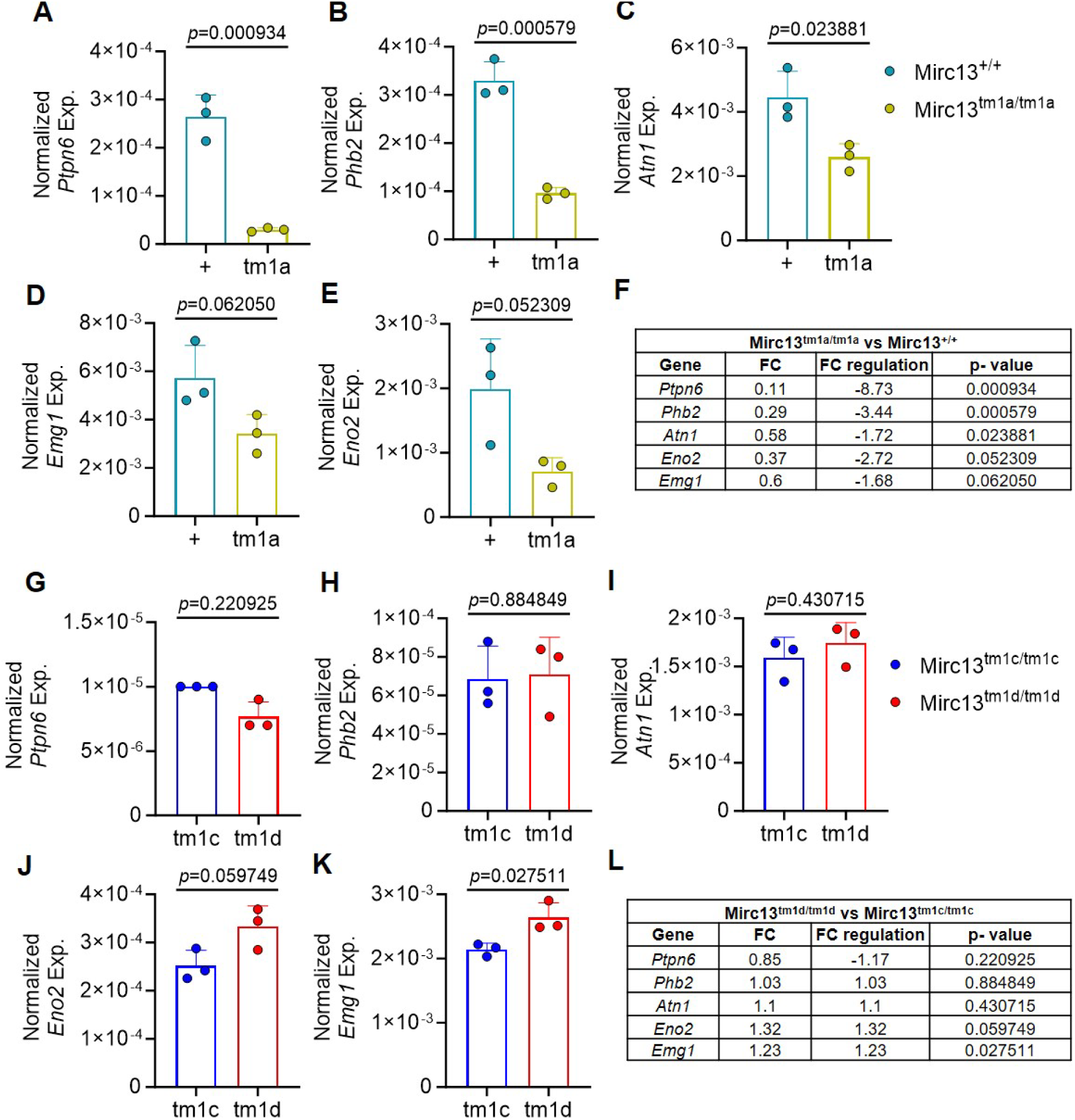
Expression analysis of Mirc13 neighboring genes. Total RNA was isolated from olfactory bulb (OB) of Mirc13^tm1a/tm1a^ and Mirc13^tm1d/tm1d^ mice and corresponding negative littermate (Mirc13^+/+^ and Mirc13^tm1c/tm1c^). Expression of *Ptpn6* (**A** and **G**), *Phb2* (**B** and **H**), *Atn1* (**C** and **I**), *Eno2* (**D** and **J**), and *Emg1* (**E** and **K**) were assessed by RT-qPCR. Data are presented as mean ± SD (*p < 0.05 vs. *Mirc13^+/+^* control. Light blue column denotes RNA obtained from *Mirc13^+/+^*; Yellow column denotes RNA obtained from *Mirc13^tm1a/tm1a^*; dark blue column denotes RNA from *Mirc13^tm1c/tm1c^* and red column denotes RNA isolated from Mirc13^tm1d/tm1d^ mice. Detail Fold Change (FC) regulation of neighboring genes (**F** & **L**). (n=3 per group, F=2 & M=1, 3-4 Week) each sample run in qPCR with technical duplicate.

### Expression Analysis of miR-141-3p and miR-200c-3p in Mirc13^tm1d/tm1d^ mice

We further characterized expression of *miR-141-3p* and *miR-200c-3p* in *Mirc13^tm1d/tm1d^* and their corresponding *Mirc13^tm1c/tm1c^*mice in the brain. We isolated total RNA from different regions of the brain, including olfactory bulbs (OB), frontal cortex (FC), parietal cortex (PC), hippocampus (HP), and cerebellum (CB), from mice of both genotypes. Our results showed both *miR-141-3p* and *miR-200c-3p* were highly expressed in the OB (ΔCT < 5; it is to be noted that lower ΔCT indicate higher expression), while their expression in other regions was very low (ΔCT > 10) (Fig 5A and B). Notably, the OB of *Mirc13^tm1d/tm1d^* mice showed no detectable expression of *miR-141-3p* by RNA-FISH (Fig 5D). This finding was further confirmed by RT-qPCR, which demonstrated a marked reduction of both miRNAs (*miR-141-3p* & *miR-200c-3p*) in the OB almost negligible (Fig 5C and C) of *Mirc13^tm1d/tm1d^* mice as compared to *Mirc13^tm1c/tm1c^*mice. Additionally, our data also showed significantly decreased expression of miR-141-3p in CB and miR-200c in the HP (Fig 5F), along decreased but not significant in FC, PC, HP (*miR-141-3p*) and FC, PC, CB (*miR-200c-3p*) (Fig 5F) of *Mirc13^tm1d/tm1d^* mice as compared to *Mirc13^tm1c/tm1c^* mice.

**Fig. 5.**
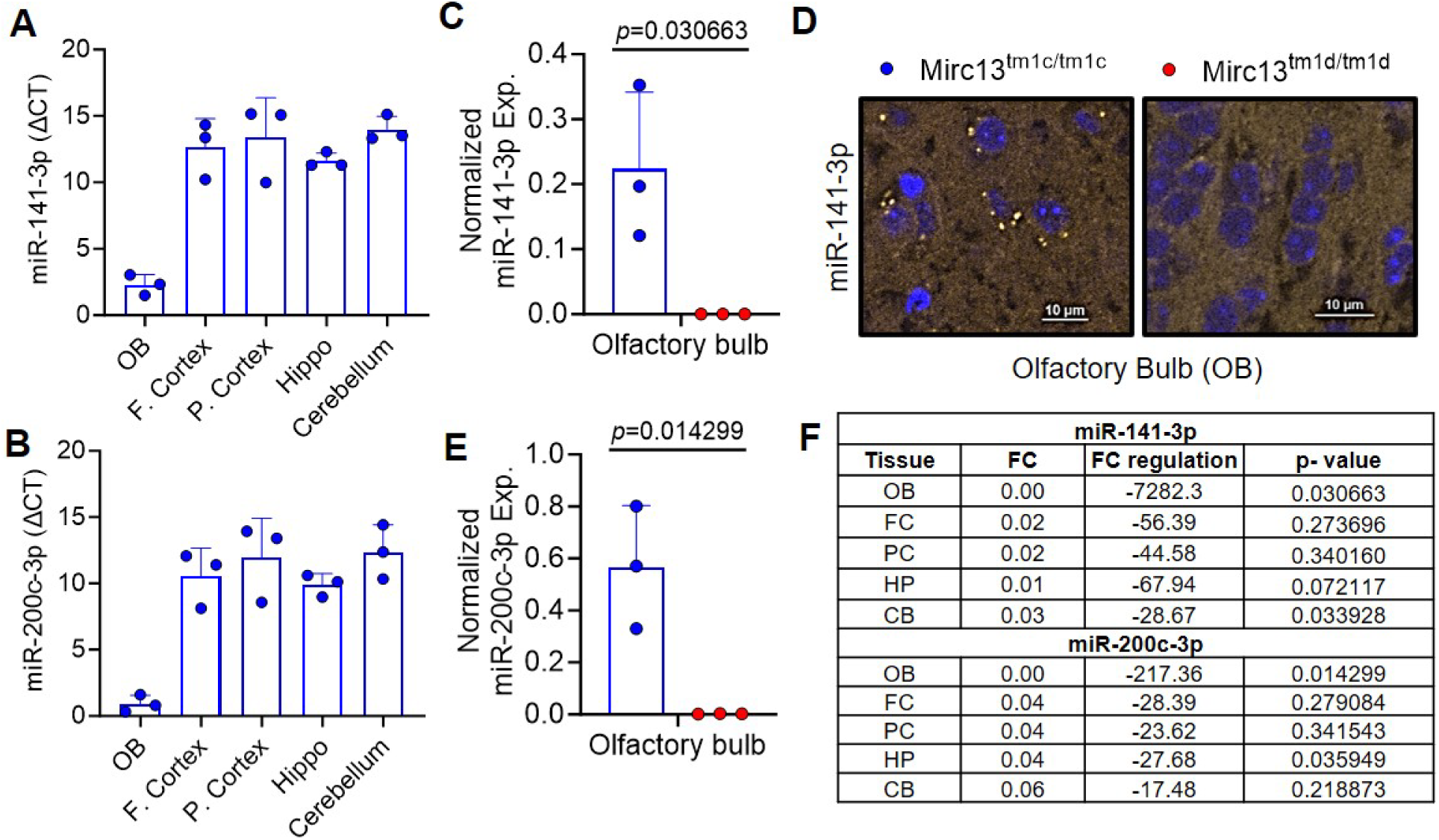
Expression analysis of miR-141-3p and miR-200c-3p in *Mirc13^tm1c/tm1c^* and *Mirc13^tm1d/tm1d^* mice. Total RNA was isolated from olfactory bulb (OB), frontal cortex (FC), parietal cortex (PC), hippocampus (HP), and cerebellum (CB) of *Mirc13^tm1c/tm1c^* and *Mirc13^tm1d/tm1d^* mice. Basal expression levels of *miR-141-3p* (**A**) and *miR-200c-3p* (**B**) were assessed by RT-qPCR using RNA isolated from *Mirc13^tm1c/tm1c^* mice. Total RNAs isolated from OB (**C** and **E**); FC; PC; HP; and CB (**F**) were examined for the normalized level of miR-141-3p and miR-200c-3p, respectively, by RT-qPCR from *Mirc13^tm1c/tm1c^* (dark blue column) and *Mirc13^tm1d/tm1d^*(red column) mice. (**D**) RNA-FISH was performed to detect expression of miR-141-3p in OB of *Mirc13^tm1c/tm1c^* and *Mirc13^tm1d/tm1d^* mice. Data are presented as mean ± SD (*p < 0.05 vs. *Mirc13^tm1c/tm1c^* control). (n=3 per group, F=2 & M=1, 3-4 Week) each sample run in qPCR with technical duplicate.

## DISCUSSION

Generation of the *Mirc13^tm1a^* line by Park et al. (2012) provided the first genetically engineered model for the miR-141/200c cluster. In their design, En2 splice acceptor and IRES sequence were placed 5’ of a lacZ reporter, followed with SV40 polyadenylation signal sequence. This design, which is similar to the gene trap design (Skarnes, et al. 1992), was aimed to reveal spatial expression of the disrupted endogenous *GM15884* as well as the miR-141/200c cluster located in intron 1 of *GM15884*. The current lacZ–polyA cassette was designed to disrupt *GM15884* transcription and substantially reduce downstream expression of the *Mirc13* locus. However, as reported for other knockout-first alleles, exogenous polyadenylation sequences may not invariably produce complete transcriptional termination because residual transcriptional read-through can occur (Lindner, et al. 2021). Therefore, although our expression analyses demonstrate marked loss of miR-141/200c expression, the present study does not directly assess the completeness of transcriptional termination. *GM15884* is a lncRNA gene predicted by bioinformatics as described in the Mouse Genome Database (Eppig, et al. 2005) (https://www.informatics.jax.org/marker/key/326668). We performed a blast search of the mouse Expressed Taq Sequence database (dbEST) (Boguski, et al. 1993), and identified a mouse cDNA clone, BB612857, of 646 bp with its 5’ sequence identical to predicted exon 1 and 2 of *GM15884*. Furthermore, the rest of BB612857 3’-downstream sequence is identical to predicted *GM15884* intron 2 sequence. It is well established that EST are derived from transcribed genes (the transcriptome) and dbEST was used to identify transcribed sequences without requiring a fully sequenced genome (Boguski, et al. 1993). Identification of BB612857 confirms the identity of *GM15884*. This result, together with the tissue-specific lacZ expression, suggests that expression of *GM15884/BB612857*, as well as *Mirc13*, is under the control of a tissue-specific Pol II promoter, and our lacZ expression observed in the olfactory bulb and lung in the mice corroborated with the finding of Park et al. (2012).

It has been reported that expression of miR-141/200c is closely associated with transcriptional read-through (TRT) bypassing the polyadenylation signal sequence of the 5’ upstream gene, *ptpn6*, or via DNA looping using two human cancer cell lines, SKOV3 and OVCA433 as well as established human and mouse fibroblast cell lines (Batista, et al. 2016). Moreover, high levels of miR-141/200c were detected under oxidative stress, while only low levels were observed under normal cell culture conditions. We have not examined whether TRT or DNA looping play any role in miR-141/200c in mice in our present studies. Nevertheless, expression of lacZ could potentially help reveal endogenous spatial expression of miR-141/200c via TRT.

Since the *Mirc13^tm1a^* mouse line was original generated by homologous recombination using ES cells, A β-actin-neo selectable marker was included in the original targeting vector and was retained in the *Mirc13^tm1a^* mouse line. Previous studies have demonstrated that promoter-containing selection cassettes can alter the expression of neighboring genes through transcriptional interference and local chromatin effects (Olson, et al. 1996; Pham, et al. 1996; West, et al. 2016). In addition, Batista et al. (2016) showed that transcription of the miR-141/200c locus can depend on transcriptional read-through and higher-order chromatin interactions, raising the possibility that the exogenous polyadenylation sequences introduced into the knockout-first allele could also disrupt local transcriptional architecture. These mechanisms are considered further below in relation to the altered neighboring-gene expression observed in the Mirc13^tm1a^ allele may result from one or more non-mutually exclusive mechanisms to be tested in future studies. Park et al (2012) explicitly recommended removal of Neo cassette prior to subsequent use of the mouse line. The Neo cassette together with the lacZ sequence were flanked by FRT sites and could be excised easily by breeding the *Mirc13^tm1a^* mice with FLP mice. Additional methods by which one could excise the FRT flanked lacZ/Neo cassette is either by electroporation of Flpo mRNA into one-cell embryos (Hashimoto and Takemoto 2015) or infection of one-cell embryos with recombinant adeno-associated viruses expressing Flpo (Nickl, et al. 2024), followed by transferring manipulated embryos into pseudopregnant females for subsequent development. Overall, it is important to generate *Mirc13^tm1c^* mice containing functional floxed *Mirc13* allele prior to using Cre recombinase for either tissue-specific or global deletion of the miR-141/200c cluster in order to avoid interference from the strong b-actin promoter neo elements, including unanticipated knockout of endogenous *GM15884* and altering the expression of neighboring genes (McBurney, et al. 2002; Branda and Dymecki 2004). Our findings are consistent with a substantial body of literature demonstrating that retained selection cassettes can influence the expression of neighboring genes. Early studies by Pham et al. (1996) and Olson et al. (1996) showed that promoter-driven neo selection cassettes can disrupt local gene regulation and generate unintended phenotypes through long-range effects on adjacent loci (Olson, et al. 1996; Pham, et al. 1996). Subsequent analyses of large-scale gene-targeting resources further highlighted retained selection cassettes as an important source of confounding transcriptional effects (Paranjape, et al. 2012). West et al. (2016) systematically analyzed targeted mouse alleles and demonstrated that heterologous promoter-driven neo cassettes frequently dysregulate neighboring genes, leading the authors to recommend careful interpretation of knockout-first alleles and consideration of cassette removal whenever possible (West, et al. 2016). Similarly, Pan et al. reported that insertion of a knockout-first cassette in the Ampd1 locus caused neonatal lethality through disruption of neighboring-gene expression (TW and Girke 2016), while Jin et al. confirmed that retention of neo-containing cassettes can produce off-target transcriptional effects in targeted mouse models (Jin, et al. 2020). Collectively, these studies underscore the importance of removing selection cassettes after gene targeting and support our observation that retention of the knockout-first cassette can alter expression of genes neighboring the Mirc13 locus.

Our neighboring-gene expression analyses demonstrate that retention of the knockout-first cassette is associated with reduced expression of several adjacent genes such as *Ptpn6, Phb2* and *Atn1* in the *Mirc13^tm1a/tm1a^*mice. However, because FLP-mediated excision simultaneously removes both the promoter-containing selection cassette and the exogenous polyadenylation sequences, our data does not distinguish whether these effects arise predominantly from promoter-mediated transcriptional interference, disruption of transcriptional read-through, altered local chromatin organization, or a combination of these mechanisms. Regardless of the underlying mechanism, our findings demonstrate that retention of the knockout-first cassette can confound interpretation of this allele and support removal of the cassette before Cre-mediated recombination. The genes *Ptpn6, Phb2,* and *Atn1* each play critical roles in brain function. *Ptpn6* regulates microglial signaling and helps restrain neuroinflammation; its loss can drive excessive immune activation and neuronal injury (He, et al. 2023). *Phb2* maintains mitochondrial integrity and neuronal survival, so its altered expression can impair energy metabolism and cognitive functions (Xu, et al. 2014). *Atn1* acts as a transcriptional co-regulator in neuronal development and disruption can lead to abnormal differentiation, connectivity, or neurodegeneration (Whitton, et al. 1993). Thus, downregulation of these genes as shown in *Mirc13^tm1a/tm1a^* mice could have significant consequences for neuronal health and brain function. There is no change in expression of these genes in *Mirc13^tm1c/tm1c^* mice. This strategy minimizes confounding effects associated with retention of the knockout-first cassette and permits accurate interpretation of miR-141/200c loss-of-function phenotypes in *Mirc13* conditional knockout mice.

Despite the clear breeding plan recommended by Park et al (Park, et al. 2012), several subsequent studies did not adopt the necessary breeding plan. *Mirc13^tm1a^* mice, which contain lacZ/β-actin-neo cassettes, were treated as *Mirc13* cKO animals and bred them directly with Cre deleters (Guan, et al. 2018; Hoefert, et al. 2018; Ji, et al. 2018; Liu, et al. 2018; Wu, et al. 2019; Xu, et al. 2025). However, *Mirc13^tm1a^* mice contain three colinear loxP sequences in the same allele (Fig 1A). In the presence of Cre, any pair of these three sites can recombine, so more than one excision product is possible (Nagy 2000; Yu, et al. 2000; Xu, et al. 2001; Branda and Dymecki 2004). The relative frequency of these products is not fixed: efficient germline deleters often favour excision of the largest loxP interval, and the outcome depends on the Cre line, expression level and timing, and transgene design. We therefore do not claim that direct Cre crossing invariably yields a heterogeneous mixture; rather, the identity of each allele must be established empirically. Figure 6 shows the possible configurations together with the primer positions used to distinguish them. As shown in Fig 6, this could generate three different genotypes dependent on the different pairings of loxP sequences: A). excision of the β-actin-Neo cassette alone; B) excision of *Mirc13* alone; and C) excision of β-actin-Neo cassette and *Mirc13* together. We have included the corresponding positions of all primers used in genotyping in Figure 1 and 6 to help identify different alleles. Nevertheless, in all of these cases, both the endogenous *GM15884* and *Mirc13* will not express, and it is still not possible to decipher the precise functional role of miR-141/200c. We used Rosa26-FLPo mice to overcome limitations associated with low FLP recombinase expression(Raymond and Soriano 2007; Wu, et al. 2009; Kranz, et al. 2010). Although FLPe is commonly used for excision, we selected FLPo because its codon optimization improves FLP expression and recombination efficiency in mammalian systems, reducing incomplete excision and mosaicism. In contrast, FLPe lacks mammalian codon optimization, and reduced FLP expression may lead to inefficient β-actin-neo cassette excision at the one-cell stage and mosaic founders. While β-actin-Flpe has been used for Mirc13^tm1a allele conversion, Rosa26-FLPo provides a more reliable strategy for generating uniformly recombined alleles (Tran, et al. 2017; Mostofa, et al. 2022; Tran, et al. 2024). The relative frequency of these products is not fixed: it depends on the Cre line, the level and timing of Cre expression, and transgene design, and efficient germline deleters often favour excision of the largest loxP interval. The identity of each allele must therefore be established empirically rather than assumed. Importantly, we did not observe evidence of mosaic recombination or incomplete cassette excision in the Mirc13 allele examined in this study. We acknowledge that recombination efficiency can vary depending on the recombinase line and target locus; however, appropriate breeding, segregation of the recombinase allele, and routine genotype verification are sufficient to identify animals carrying fully recombined alleles before experimental use. Our breeding plan, as shown in Fig 1, is to remove the lacZ/neo cassettes using ROSA26-FLPo mice, generating *Mirc^tm1c^*, which is a functional *Mirc13* floxed, allele prior to using Cre mice, such as the germline *Hprt^tm1(Cre)^* mice, to achieve global deletion of the *Mirc13* allele. These steps ensure that any observed phenotype will be attributed directly to loss of the miR-141/200c cluster, without any confounding contributions from transcriptional silencing or residual neomycin cassette effects. The scope of this recommendation warrants comment. The mechanism is not miRNA-specific, but its magnitude varies with local genomic architecture, so our findings should not be read as a blanket statement about all knockout-first alleles. This is relevant to the standard tm1a-to-tm1b conversion, in which Cre removes the β-actin-neo cassette together with the critical exon while the En2 splice acceptor-IRES-lacZ-polyA trap is retained by design: the heterologous promoter is eliminated, but the splice acceptor and exogenous polyadenylation signal are not, so the potential for disrupted read-through persists. Because tm1a carries three colinear loxP sites, tm1b is also only one of several possible recombination products and must be confirmed by genotyping rather than assumed (Fig. 6). We therefore recommend that, wherever a cassette or trap is retained by design, expression across the surrounding interval be measured directly or the phenotype cross-validated in cassette-free animals at the same locus, as done here for *Mirc13^tm1c^* and *Mirc13^tm1d^*.

**Fig. 6.**
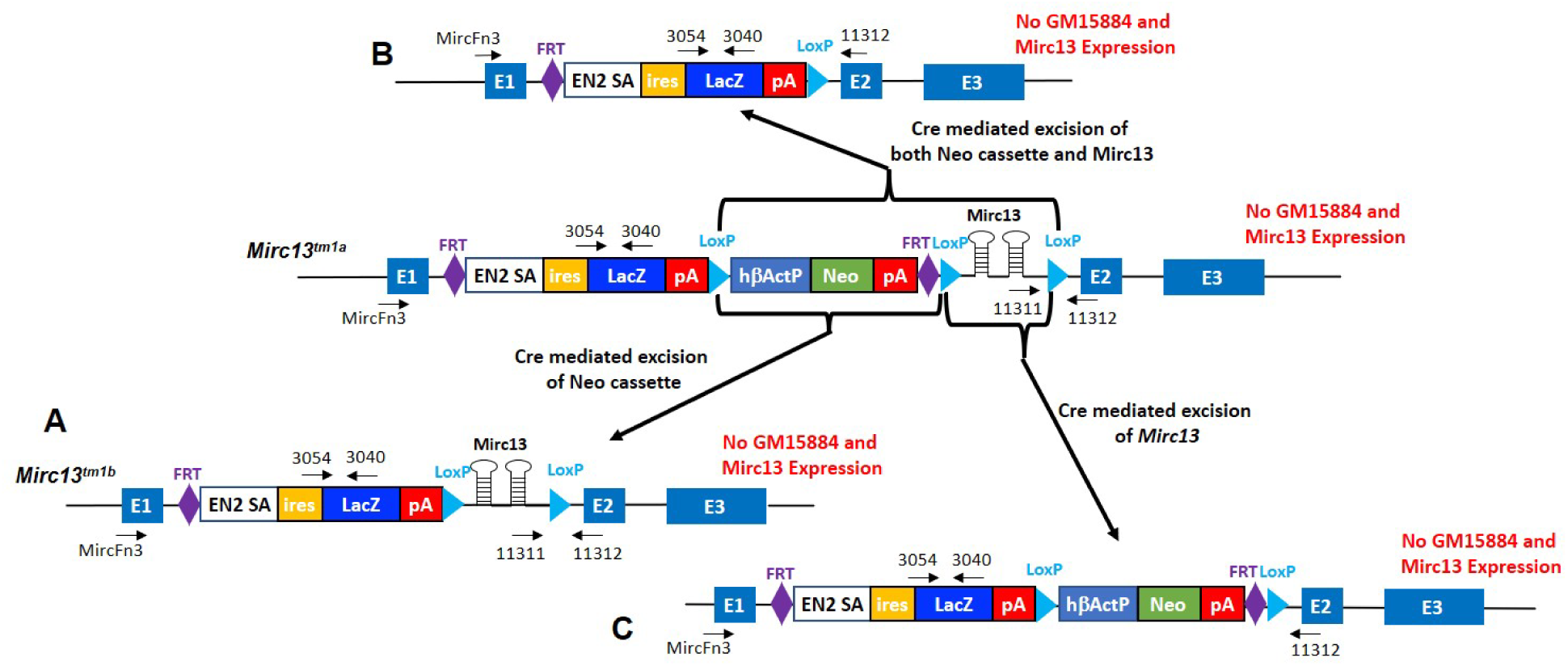
Breeding Mirc13^tm1a^ mice containing three colinear loxP sites with Cre mice could lead to three different genotypes. A Excision of the β-actin-Neo cassette only to generate the tm1b allele with retention of the lacZ reporter; B excision of β-actin-Neo cassette together with Mirc13 sequence; and C deletion of Mirc13 cluster only. In all three different genotypes, expressions of both GM15884 and Mirc13 are disrupted due to the presence of the En2-IRES-lacZ and polyA sequence.

This breeding plan is not merely technical but critical for precise evaluation of the miR-141/200c functional roles. Studies of the miR-200 family—including their roles in late postnatal neurogenesis (Beclin, et al. 2016), stem cell differentiation (Zhang, et al. 2015), and cancer biology (Chen, et al. 2013; Jin, et al. 2017; Tseng, et al. 2017) rely strongly on precise gene dosage and context. Using animals with imprecise genotypes will only hamper validities of the data leading to spurious conclusion and discrepancy among different reports. By adhering to the breeding plan according to the design of the original targeting vector, our work provides a framework to generate unambiguous global and functional conditional knockouts of the miR-141/200c cluster. We also acknowledge several limitations. First, although our results consistently show altered neighboring gene expression in the knockout-first allele, the small sample size (n = 3 per genotype) limits the precision of effect-size estimation and warrants cautious interpretation. Second, because FLP-mediated excision removes the beta-actin promoter and the exogenous polyadenylation signals simultaneously, our design cannot attribute the effect to either mechanism; strand-specific RNA-seq or 3′ RACE would be required to do so. Third, expression profiling was restricted to the olfactory bulb, the tissue with the highest miR-141/200c expression, so tissue-specific differences elsewhere cannot be excluded. Fourth, the cohorts were not powered to assess sex as a biological variable.

## Author contributions

RV conceptualized the project. SKY, RS, RV and SPY wrote the manuscript. MKC, RS, SKY and KL performed the experiments. RV and SPY supervised the work. RV secured the funding for the project. All the authors approved the work.

## Ethics declarations

### Ethical Approval

All experiments were approved and conducted in compliance with the protocols approved (Number: AP-201043-0926) by the Institutional Animal Care and Use Committee (IACUC) at UConn Health. Animal data was reported following the ARRIVE Guidelines.

### Data Availability

All data supporting the findings are included in the article, that the validated *Mirc13*^tm1c^ and *Mirc13*^tm1d^ lines will be deposited in a public repository (JAX/MMRRC) upon acceptance.

